# Stereoinversion via alcohol dehydrogenases enables complete catabolism of β-1-type lignin-derived aromatic isomers

**DOI:** 10.1101/2023.01.24.525472

**Authors:** Ryo Kato, Kodai Maekawa, Shota Kobayashi, Shojiro Hishiyama, Rui Katahira, Miki Nambo, Yudai Higuchi, Eugene Kuatsjah, Gregg T. Beckham, Naofumi Kamimura, Eiji Masai

## Abstract

*Sphingobium* sp. SYK-6 is an efficient aromatic catabolic bacterium that can consume all four stereoisomers of 1,2-diguaiacylpropane-1,3-diol (DGPD), which is a ring-opened β-1-type dimer. Recently, LdpA-mediated catabolism of *erythro*-DGPD was reported in SYK-6, but the catabolic pathway for *threo*-DGPD was heretofore unknown. Here we elucidated the catabolism of *threo*-DGPD, which proceeds through conversion to *erythro*-DGPD. When *threo*-DGPD was incubated with SYK-6, the Cα alcohol groups of *threo*-DGPD (DGPD I and II) were initially oxidized to produce the Cα carbonyl form (DGPD-keto I and II). This initial oxidation step is catalyzed by Cα-dehydrogenases, which belong to the short-chain dehydrogenase/reductase (SDR) family and are involved in the catabolism of β-O-4-type dimers. Analysis of seven candidate genes revealed that NAD^+^-dependent LigD and LigL are mainly involved in the conversion of DGPD I and II, respectively. Next, we found that DGPD-keto I and II were reduced to *erythro*-DGPD (DGPD III and IV) in the presence of NADPH. Genes involved in this reduction were sought from Cα-dehydrogenase and *ldpA*-neighboring SDR genes. The gene products of SLG_12690 (*ldpC*) and SLG_12640 (*ldpB*) catalyzed the NADPH-dependent conversion of DGPD-keto I to DGPD III and DGPD-keto II to DGPD IV, respectively. Mutational analysis further indicated that *ldpC* and *ldpB* are predominantly involved in the reduction of DGPD-keto. Together, these results demonstrate that SYK-6 harbors a comprehensive catabolic enzyme system to utilize all four β-1-type stereoisomers through successive oxidation and reduction reactions of the Cα alcohol group of *threo*-DGPD with a net stereoinversion using multiple dehydrogenases.

**IMPORTANCE:** In many catalytic depolymerization processes of lignin polymers, aryl–ether bonds are selectively cleaved, leaving carbon–carbon bonds between aromatic units intact, including dimers and oligomers with β-1 linkages. Therefore, elucidating the catabolic system of β-1-type lignin-derived compounds will aid in the establishment of biological funneling of heterologous lignin-derived aromatic compounds to value-added products. In this work, we found that *threo*-DGPD was converted by successive stereoselective oxidation and reduction at the Cα-position by multiple alcohol dehydrogenases to *erythro*-DGPD, which is further catabolized. This system is very similar to that developed to obtain enantiopure alcohols from racemic alcohols by artificially combining two enantiocomplementary alcohol dehydrogenases. The results presented here demonstrate that SYK-6 has evolved to catabolize all four stereoisomers of DGPD by incorporating this stereoinversion system into its native β-1-type dimer catabolic system.

## INTRODUCTION

Lignin, a heteropolymer composed of phenylpropanoids, is a major component of plant cell walls, accounting for up to ~30% of the plant biomass (1). Since lignin is the most abundant bio-based aromatic resource, second only to cellulose, lignin valorization is a critical need to enable a lignocellulosic bioeconomy. In recent years, there has been substantial interest in processes that produce value-added products from lignin by combining catalytic depolymerization of lignin with biological funneling, or convergent microbial conversion of mixed compounds to a single target product. These processes have enabled the laboratory-scale production of multiple value-added products (2–9). Given this intense level of interest in this topic, it is becoming increasingly important to elucidate the bacterial catabolic systems for the conversion of lignin-derived aromatic compounds with high efficiency and yield (2–10).

Lignin contains various C–O–C and C–C bonds such as β-O-4, β-5, 5-5, β-1, and β-β linkages formed between phenylpropane units during lignin biosynthesis (11, 12). In addition, despite having many asymmetric carbons, lignin is optically inactive, and thus is considered to be a racemate containing equal amounts of optical isomers (13). Among the lignin intermolecular linkages, the β-1 linkage exists in native lignin as the spirodienone structure. Nevertheless, spirodienone is unstable and readily undergoes hydrolysis under mild acidic conditions, which releases glyceraldehyde-2-aryl ether to produce the chemically-stable diarylpropane (14). The β-1 linkage is present in 1–9% of lignin, and it typically remains intact, along with other C–C bonded dimers and oligomers that are obtained in catalytic depolymerization processes that selectively cleave the β-O-4 linkage (15–19). Therefore, to enable higher product yields from lignin-derived compounds, it would be beneficial to harness the bacterial catabolic system for β-1-type lignin-derived compounds.

*Sphingobium* sp. SYK-6 (hereafter SYK-6) is a Gram-negative aerobic bacterium belonging to the alphaproteobacteria class and is the best-characterized bacterium capable of degrading lignin-derived aromatic dimers and monomers (20–23). To date, the catabolic system of β-O-4, β-5, and 5-5 dimers has been largely elucidated, but until recently, almost nothing was known about the catabolic system of β-1 dimers. A β-1-type dimer, 1,2-diguaiacylpropane-1,3-diol (DGPD), has *threo* and *erythro* diastereomers, each with two enantiomers. (**Fig. 1A**). The bacterial DGPD catabolic system was first reported about 30 years ago in *Sphingomonas paucimobilis* TMY1009. In *S. paucimobilis* TMY1009, the *erythro*-diastereomer of DGPD (*erythro*-DGPD) undergoes Cγ deformylation accompanied with dehydroxylation at the Cα position to form a stilbene-type compound [1,2-bis(4-hydroxy-3-methoxyphenyl)ethylene); DGPD-S], which is then degraded to two vanillin molecules via Cα-Cβ cleavage by lignostilbene α,β-dioxygenase (24–27).

**FIG 1.**
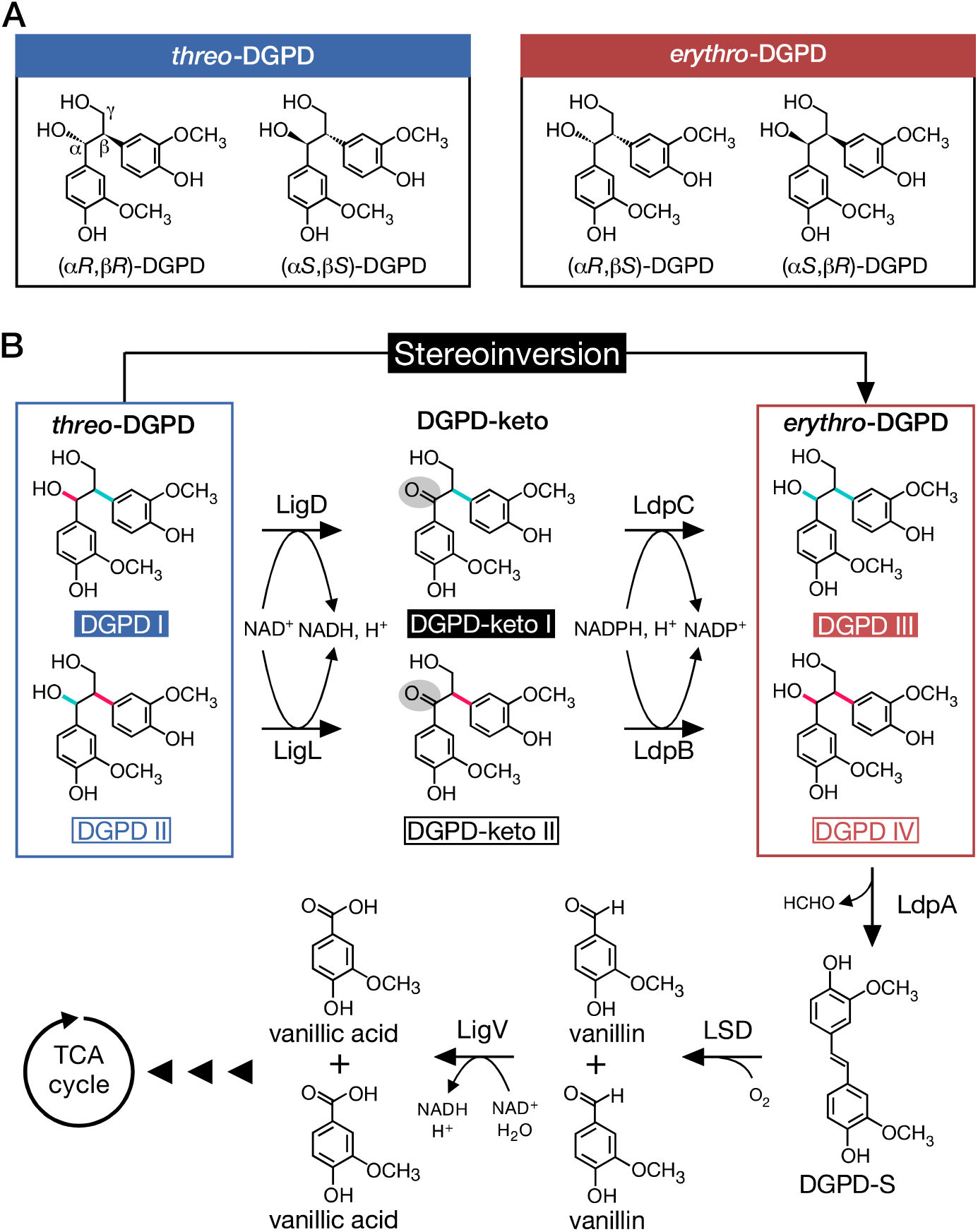
Catabolic pathway of four stereoisomers of a β-1-type lignin-derived aromatic dimer in *Sphingobium* sp. strain SYK-6. (A) Structure of stereoisomers of 1,2-diguaiacylpropane-1,3-diol (DGPD). (B) Catabolic pathway of *threo*-DGPD. Since the absolute configuration of each isomer is unknown, each isomer is distinguished by changing the color of the bonds from the asymmetric carbon at the α-position to the hydroxy group and the asymmetric carbon at the β-position to the 4-hydroxy-3-methoxyphenyl group. Enzymes: LigD, LigL, LdpC, and LdpB, short-chain dehydrogenase/reductases; LdpA, *erythro*-DGPD Cγ-formaldehyde lyase; LSD, lignostilbene α,β-dioxygenase; LigV, vanillin dehydrogenase.

The *erythro*-DGPD converting enzyme was purified in this study, but the enzyme and gene sequences were not obtained. In 2021, the *erythro*-DGPD Cγ-formaldehyde lyase gene (originally dubbed *lsdE*, but later renamed *ldpA*) was identified in *Novosphingobium aromaticivorans* DSM 12444 using randomly barcoded transposon insertion sequencing analysis (28). Most recently, the ortholog of Cγ-formaldehyde lyase gene of *N. aromaticivorans* DSM 12444 was found to be responsible for *erythro-DGPD* conversion in SYK-6, and the structures of LdpA of SYK-6 and *N. aromaticivorans* DSM 12444 were determined and the reaction mechanism was proposed (29). Interestingly, LdpA exhibited conversion activity specifically for *erythro*-DGPD, but not for the *threo*-diastereomer of DGPD (*threo*-DGPD) mainly due to the inability to form proper hydrogen bonds between the substrate and enzyme.

In this study, we found that the two enantiomers of *threo*-DGPD are stereoselectively oxidized to the two enantiomers of the Cα carbonyl form, DGPD-keto, which is further stereoselectively reduced to the two enantiomers of *erythro*-DGPD that are catabolized via the LdpA-mediated reaction. Furthermore, the enzyme genes involved in these conversions were identified, characterized, and the entire SYK-6 β-1 dimer catabolic system was elucidated. The results revealed that SYK-6 has evolved to degrade all four stereoisomers of DGPD by incorporating a net stereoinversion system.

## RESULTS

### Identification of a metabolite of *threo*-DGPD in *Sphingobium* sp. SYK-6

Our previous study showed that SYK-6 can grow on *erythro*-DGPD as a sole carbon and energy source but cannot grow on *threo*-DGPD (29). To determine the ability of SYK-6 to convert *threo-*DGPD, resting cells of SYK-6 grown in lysogeny broth (LB) were incubated with 100 μM *threo-*DGPD for 24 h, and the reaction mixture was analyzed using high-performance liquid chromatography (HPLC). After the incubation, *threo*-DGPD disappeared almost completely. To identify metabolites of *threo*-DGPD, cell extracts of SYK-6 grown in LB were reacted with 100 μM *threo*-DGPD for 6 h and analyzed using HPLC-mass spectrometry (HPLC-MS), but substrate conversion was not observed. When the same reaction was performed in the presence of NAD^+^ or NADP^+^, a marked decrease in *threo*-DGPD and formation of an unknown compound I with a retention time of 2.5 min were observed (**Fig. 2A** and **B; Fig. S1**). Comparison of retention times (**Fig. 2B** and **F**), UV-visible spectra (**Fig. 2C** and **G**), and mass spectra (**Fig. 2D** and **H**) with the authentic compound revealed that compound I is DGPD-keto, in which the Cα alcohol group of *threo*-DGPD is oxidized to a carbonyl group (**Fig. 2E**). Thus, the catabolism of *threo*-DGPD in SYK-6 appears to proceed via DGPD-keto intermediate (**Fig. 1B**).

**FIG. 2.**
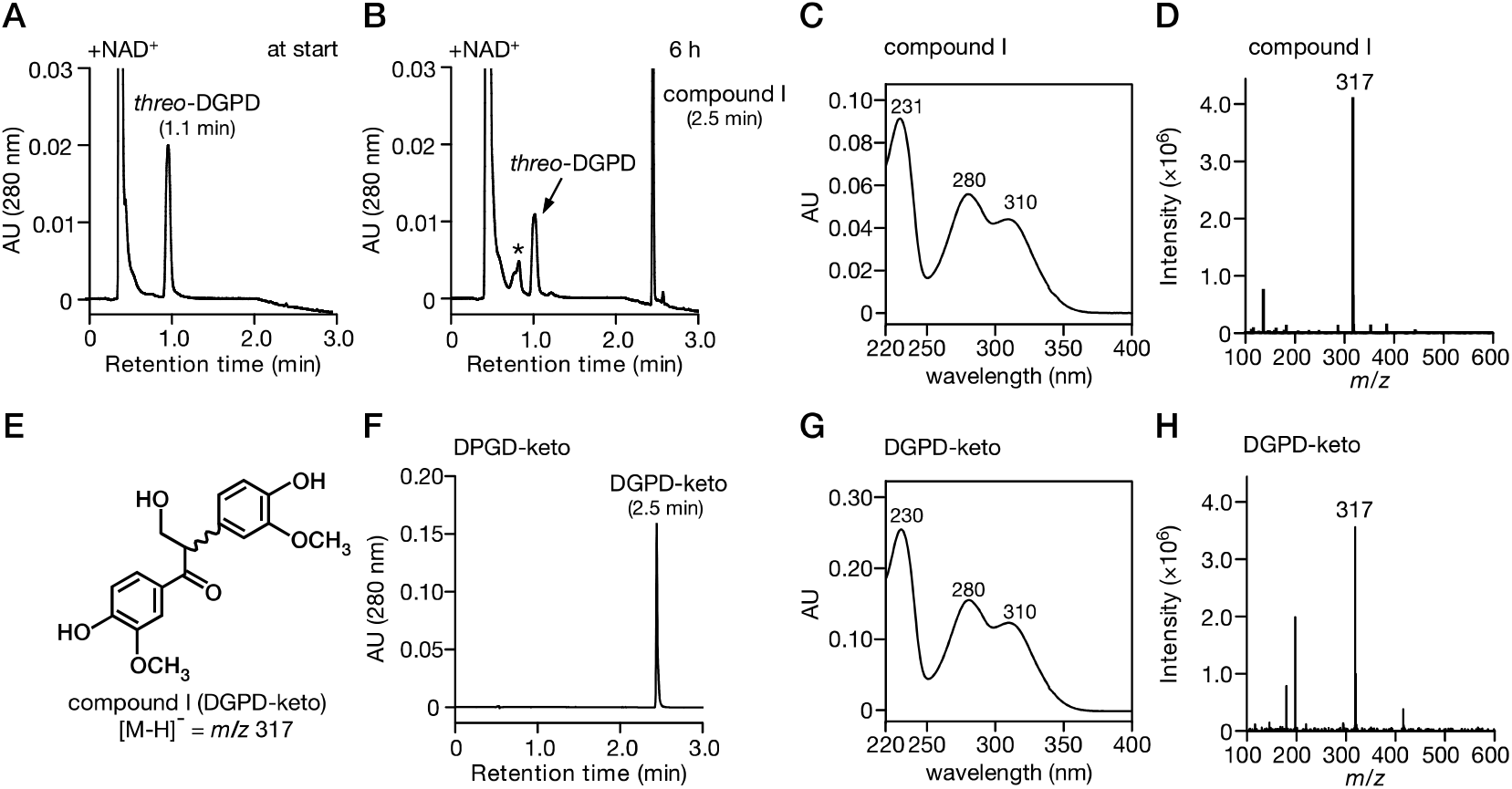
Identification of a metabolite of *threo*-DGPD. *threo*-DGPD (100 μM) was reacted with a cell extract of SYK-6 (2 mg protein/mL) in the presence of 1 mM NAD^+^. (**A**, **B**) Portions of the reaction mixtures were collected at the start and after 6 h of reaction and analyzed by HPLC-MS. *The peak with a retention time of 0.82 min appears to be an unknown compound derived from the cells. (**C**, **D**) The UV-visible spectrum and electrospray ionization (ESI)–MS spectrum of compound I (negative-ion mode) are shown. (**E**) Chemical structure of DGPD-keto. (**F**–**H**). HPLC chromatogram, UV-visible spectrum, and ESI-MS spectrum of authentic DGPD-keto.

### Identification of Cα-dehydrogenases with *threo*-DGPD conversion activity

Cα-dehydrogenases (LigD, LigL, and LigN) of SYK-6, which belong to the short-chain dehydrogenase/reductase (SDR) family, catalyze the oxidation of the Cα alcohol group of guaiacylglycerol-β-guaiacyl ether (GGE), a model β-O-4-type compound (**Fig. S2**). The reactions catalyzed by LigD, LigL, and LigN are essential for the catabolism of β-O-4-type compounds in SYK-6 (30). Specifically, LigD stereoselectively converts (α*R*,β*S*)-GGE and (α*R*,β*R*)-GGE, while LigL and LigN stereoselectively convert (α*S*,β*R*)-GGE and (α*S*,β*S*)-GGE. Additionally, another Cα-dehydrogenase, LigO, which is less involved in GGE catabolism but exhibits the same stereoselectivity as LigD, was found in SYK-6 (30). DGPD and GGE are similar in that both have two aromatic rings attached to 1,3-propanediol through carbon–carbon or aryl ether bonds and chirality at the Cα and Cβ (**Fig. 1B; Fig. S2**). In *S. paucimobilis* TMY1009, two purified Cα-dehydrogenases were reported to be able to oxidize the Cα alcohol group of both GGE and *erythro*-DGPD (31). Together, these observations indicate that Cα-dehydrogenases are likely involved in the oxidation of the Cα alcohol group of *threo*-DGPD.

Phylogenetic analysis based on amino acid sequence alignment of the SDR family genes in SYK-6, including the known Cα-dehydrogenase genes, revealed that SLG_07240, SLG_12640, and SLG_28340 are in the same clade as LigD, LigL, LigN, and LigO (**Fig. 3**). Here we defined the group containing these seven genes as the Cα-dehydrogenase group and examined the ability of each gene product to convert *threo*-DGPD. *ligD*, *ligO*, *ligL*, *ligN*, SLG_12640, and SLG_28340 were expressed as His-tag fusions under the control of the T7 promoter in *E. coli* (**Fig. S3**). Since SLG_07240 was not expressed in *E. coli*, it was expressed as a His-tag fusion under the control of the Q5 promoter in SYK-6 (**Fig. S3**). Each gene product was subsequently purified by Ni-affinity chromatography (**Fig. S3**). Since *threo*-DGPD contains two enantiomers, both enantiomers were separated using chiral HPLC. The peak eluting first in this analysis was designated as DGPD I, and the peak eluting later was designated as DGPD II (**Fig. 4A**). To determine whether Cα-dehydrogenases can convert *threo*-DGPD, each purified enzyme was reacted with *threo*-DGPD in the presence of NAD^+^ or NADP^+^. Chiral HPLC analysis showed that in the presence of NAD^+^, LigD, LigO, and the gene product of SLG_12640 converted DGPD I to DGPD-keto I (the later-eluting enantiomer in the authentic racemic DGPD-keto, **Fig. 4A**), while LigL and LigN converted DGPD II to DGPD-keto II (the first eluting enantiomer in the authentic racemic preparation of DGPD-keto) (**Fig. 4A**). However, the gene products of SLG_07240 and SLG_28340 failed to convert either substrate. When NADP^+^ was used, only LigL converted DGPD II, but to a lesser extent than in the presence of NAD^+^ (**Fig. S4**). The specific activities of LigD, LigO, SLG_12640, LigL, and LigN toward DGPD I and II were measured in the presence of NAD^+^ (**Table 1**). The SLG_12640 product and LigO showed the highest specific activities toward DGPD I (0.022–0.026 μmol·min^-1^·mg^-1^), while LigD had a specific activity of approximately 50-60% of them. LigL exhibited by far the highest specific activity toward DGPD II (5.2 μmol·min^-1^·mg^-1^), which was more than 200-fold higher than those of the SLG_12640 product and LigO toward DGPD I.

**FIG. 3.**
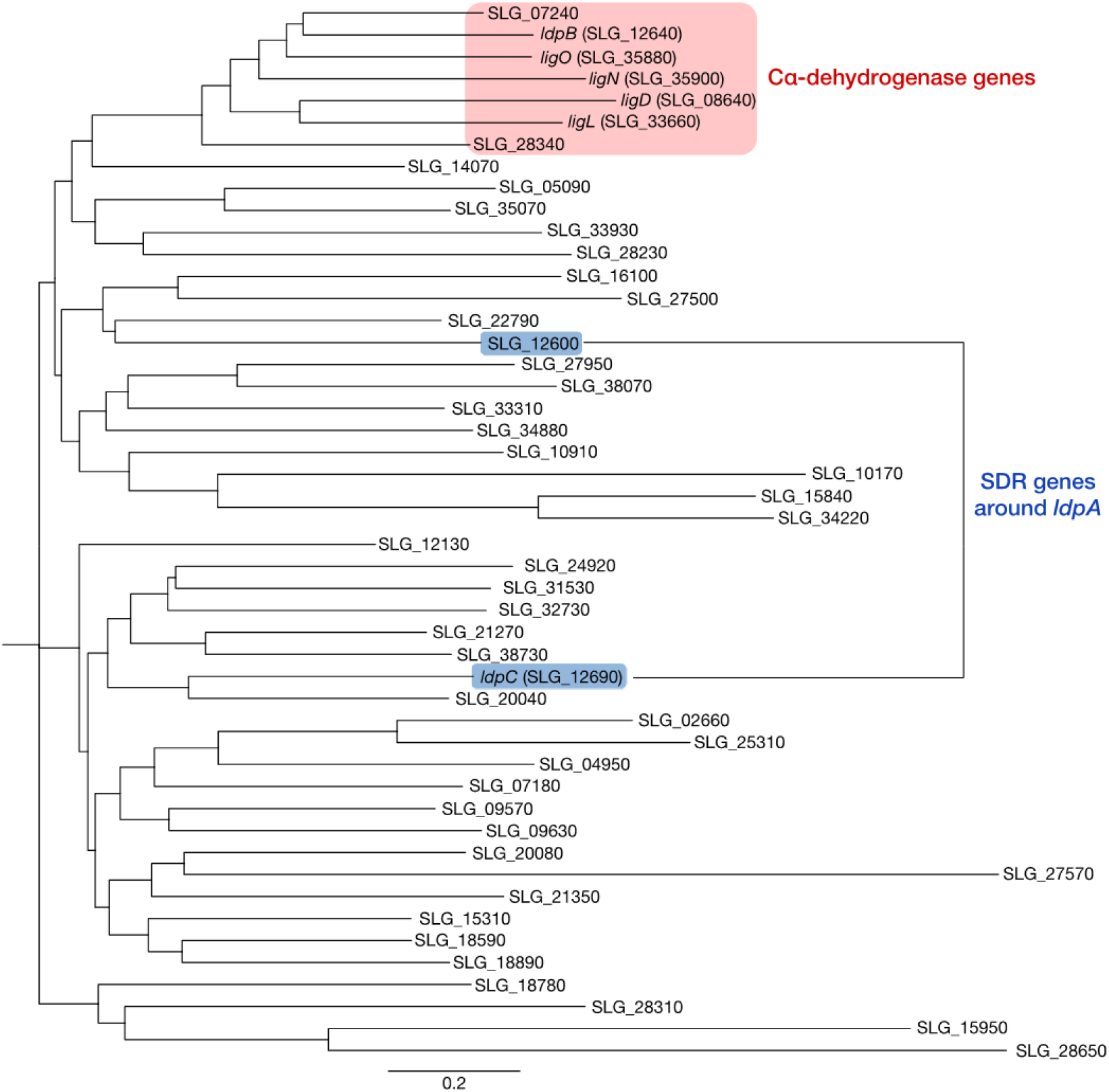
Phylogenetic tree of SDRs from SYK-6. The group containing SLG_07240, SLG_12640, and SLG_28340 with LigD, LigL, LigN, and LigO was defined as the Cα-dehydrogenase group (red background). In addition to the Cα-dehydrogenase group, SLG_12600 and SLG_12690, located around *ldpA* (blue background), were investigated in this study.

**FIG. 4.**
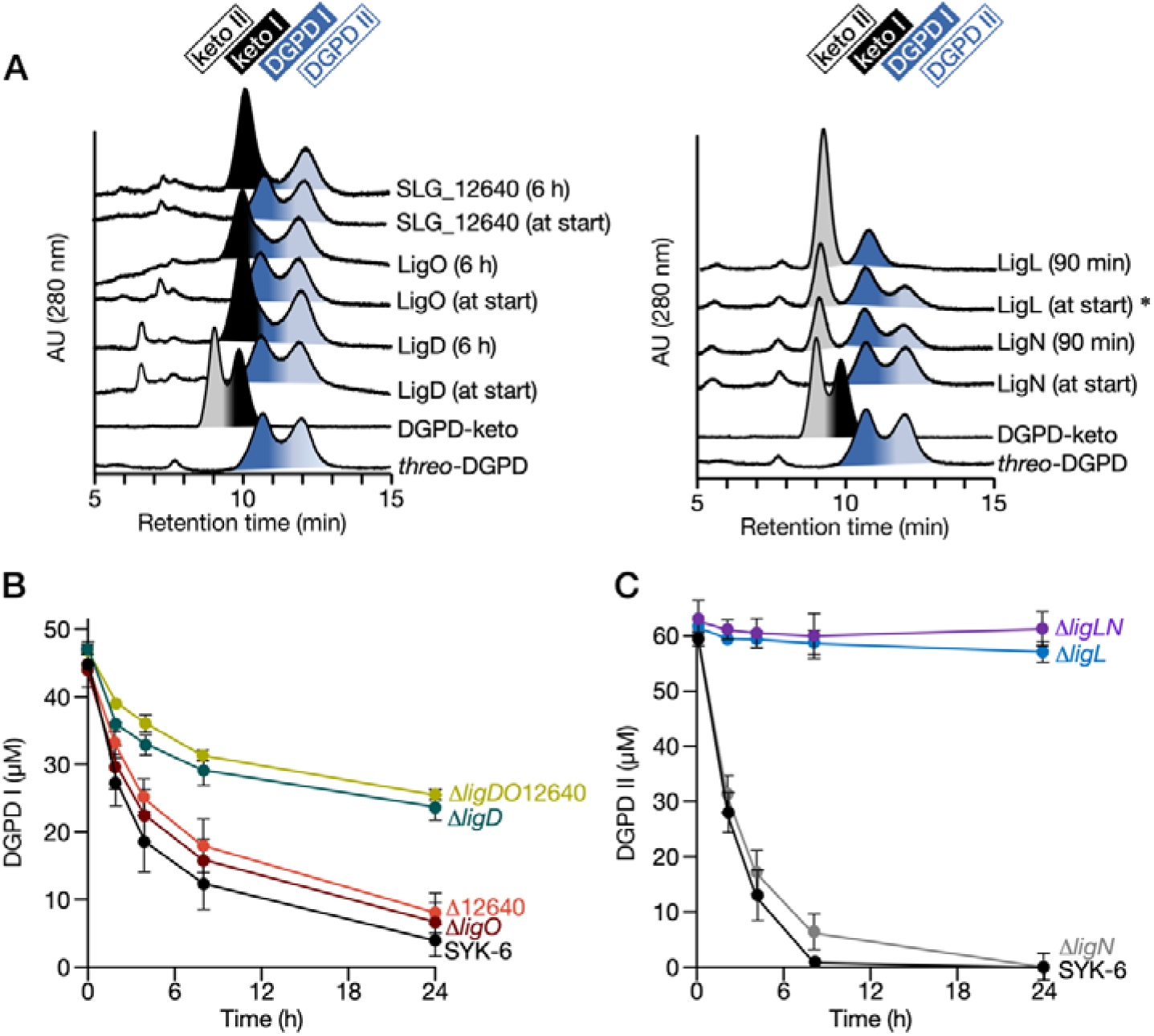
Identification of the gene products involved in *threo*-DGPD conversion in SYK-6. (**A**) Chiral HPLC analysis of the products generated from *threo*-DGPD in the reactions catalyzed by Cα-dehydrogenases. *threo*-DGPD (100 μM) was reacted with purified Cα-dehydrogenases (LigD, LigO, SLG_12640 gene product [LdpB], and LigN, 100 μg protein/mL; LigL, 1 μg protein/mL) in the presence of 1 mM NAD^+^. Portions of the reaction mixtures were collected at the start, after 90 min, or 6 h, and analyzed by chiral HPLC. *DGPD II conversion by LigL was proceeding at the start. (**B**, **C**) Conversion of 50 μM DGPD I (**B**) and II (**C**) by cells of SYK-6 and disruption mutants of the Cα-dehydrogenase genes. For DGPD I and II conversion, cells were used with an OD_600_ of 10 and 1.0, respectively. Portions of the reaction mixtures were collected and analyzed by HPLC. All experiments were performed in triplicate, and each value represents the mean ± standard deviation.

**TABLE 1.** Specific activities of Cα-dehydrogenases for DGPD I and II.

| Enzyme | DGPDI |  | DGPDI I |  |
| --- | --- | --- | --- | --- |
| | $(\mu\text{mol}\cdot\text{min}^{-1}\cdot\text{mg}^{-1})$ | | $(\mu\text{mol}\cdot\text{min}^{-1}\cdot\text{mg}^{-1})$ | |
|  | NAD<sup>+</sup> | NADP<sup>+</sup> | NAD<sup>+</sup> | NADP<sup>+</sup> |
| LigD | 0.013 ± 0.001 | ND | – | – |
| LigO | 0.022 ± 0.007 | ND | – | – |
| SLG_12640 (LdpB) | 0.026 ± 0.016 | ND | – | – |
| LigL | – | – | 5.2 ± 0.2 | ND |
| LigN | – | – | 0.070 ± 0.018 | ND |

Purified α-dehydrogenases (LigD, LigO, SLG\_12640 [LdpB], and LigN, 100 $\mu\text{g}$ protein/mL; LigL, 1 $\mu\text{g}$ protein/mL) were reacted with 50 $\mu\text{M}$ DGPDI or II in the presence of 500 $\mu\text{M}$ NAD<sup>+</sup> or NADP<sup>+</sup>. The data represent the averages ± standard deviations of three independent experiments. ND, not detected. –, not tested.

### *threo*-DGPD conversion by SYK-6 mutants of the Cα-dehydrogenase genes

To identify the genes involved in *threo*-DGPD conversion in SYK-6, we created the disruption mutants of *ligD*, *ligO*, SLG_12640, *ligL*, and *ligN* (Δ*ligD*, Δ*ligO*, Δ12640, ΔligL, Δ*ligN*); the *ligD*, *ligO*, and SLG_12640 triple disruption mutant (Δ*ligDO12640*); and the *ligL* and *ligN* double disruption mutant (Δ*ligLN*) (**Fig. S5**). Resting cells of each disruption mutant grown in LB were incubated with DGPD I or DGPD II. The amount of DGPD I converted by Δ*ligD* after 4 h of incubation was reduced to approximately 54% of that of SYK-6, while the amount of DGPD I converted by Δ*ligO* and Δ*12640*, respectively, was reduced to approximately 80% of that of SYK-6 (**Fig. 4B**). The amount of DGPD I converted by Δ*ligDO12640* was further reduced to approximately 40% of that of SYK-6. For DGPD II conversion, Δ*ligL* and Δ*ligLN* showed little conversion ability, while Δ*ligN* displayed conversion ability comparable to SYK-6 (**Fig. 4C**). Δ*ligD* and Δ*ligL* cells transformed by complementation plasmids regained the ability to convert DGPD I and II, respectively (**Fig. S6**), indicating that LigD is mainly involved in the conversion of DGPD I and LigL in the conversion of DGPD II in SYK-6.

Comparing the activity of SYK-6 resting cells toward DGPD I and II, SYK-6 showed higher activity toward DGPD II (45.8 μM·h^-1^ per cells with an OD_600_ of 1.0), which was approximately 17-fold higher than that toward DGPD I (**Fig. 4B** and **C**). In contrast, the activity of LigL toward DGPD II was 400-fold higher than that of LigD toward DGPD I. This difference is considerably greater than the difference in activity in SYK-6 (**Table 1**). The reason for this is unclear, given that LigD plays a significant role in the conversion of DGPD I. LigD is an SDR that typically forms homodimers, but it may have the potential to form a heteromer with other unidentified SDRs (32, 33). In addition, differences in *ligD* and *ligL* transcription levels may also have an effect.

Since the conversion of DGPD I and II by LigD and LigL is an oxidation reaction by alcohol dehydrogenases (ADHs), the reduction reaction (reverse reaction) may proceed using the products DGPD-keto and NADH as substrates. Therefore, LigD and LigL were reacted with DGPD-keto I and II in the presence of 1 mM NADH, respectively. When LigD reacted with DGPD-keto I, little was converted, and when LigL reacted with DGPD-keto II, only a tiny amount of DGPD was produced (**Fig. S7**). Since the activities of the reverse reaction by LigD and LigL are apparently quite low, the DGPD-keto produced is likely to proceed to the next reaction.

### Conversion of DGPD-keto I and II by SYK-6 cell extracts

To investigate the ability of SYK-6 to convert DGPD-keto I and II, resting cells (OD_600_ of 1.0) of SYK-6 grown in LB were incubated with DGPD-keto I and II for 24 h, and the reaction products were analyzed by HPLC. DGPD-keto I and II were almost fully absent after 8 h of incubation. To identify metabolites of DGPD-keto I and II, cell extracts (500 μg protein/mL) of SYK-6 grown in LB were reacted with DGPD-keto I and II for 1 h. However, no DGPD-keto I and II conversions were observed. The same reactions were performed in the presence of NADH and NADPH. As a result, no conversion of DGPD-keto I and II was observed in the presence of NADH (**Fig. S8**). In contrast, DGPD-keto I and II were converted in the presence of NADPH, and vanillic acid was detected as a product from both substrates. (**Fig. 5A** and **B**). In addition to vanillic acid, a small peak consistent with the retention time of *threo/erythro-DGPD* (1.1 min) was observed. Therefore, to clarify whether *erythro-DGPD* was generated from DGPD-keto, cell extracts of an SYK-6 disruption mutant of *ldpA* (Δ*ldpA*), which can no longer convert *erythro*-DGPD, were reacted with DGPD-keto I and II in the presence of NADPH. HPLC analysis showed a decrease in both DGPD-keto I and II, and accumulation of DGPD as judged by the retention time and UV-visible spectrum of the authentic *erythro*-DGPD (**Fig. 5C** and **D**; **Fig. S9**). Chiral HPLC analysis was performed to confirm whether the product was *erythro-DGPD;*however, the DGPD-keto I conversion product was not clearly identified because the peaks of *erythro-*DGPD and DGPD-keto I overlapped (**Fig. 5E**). In contrast, no *threo*-DGPD was generated in this reaction, thus demonstrating that the DGPD generated during the DGPD-keto I conversion was indeed the *erythro* form. With respect to DGPD-keto II conversion, the formation of *erythro*-DGPD was clearly observed (**Fig. 5F**). Together, these results indicate that *threo*-DGPD (DGPD I and II) is first oxidized to DGPD-keto I and II in SYK-6, followed by reduction to *erythro*-DGPD (DGPD III and DGPD IV) by NADPH-dependent enzymes. This fact implies that *threo*-DGPD undergoes stereoinversion of the Cα-hydroxy group to generate the *erythro* form, which is further converted by the *erythro*-DGPD-specific LdpA enzyme.

**FIG. 5.**
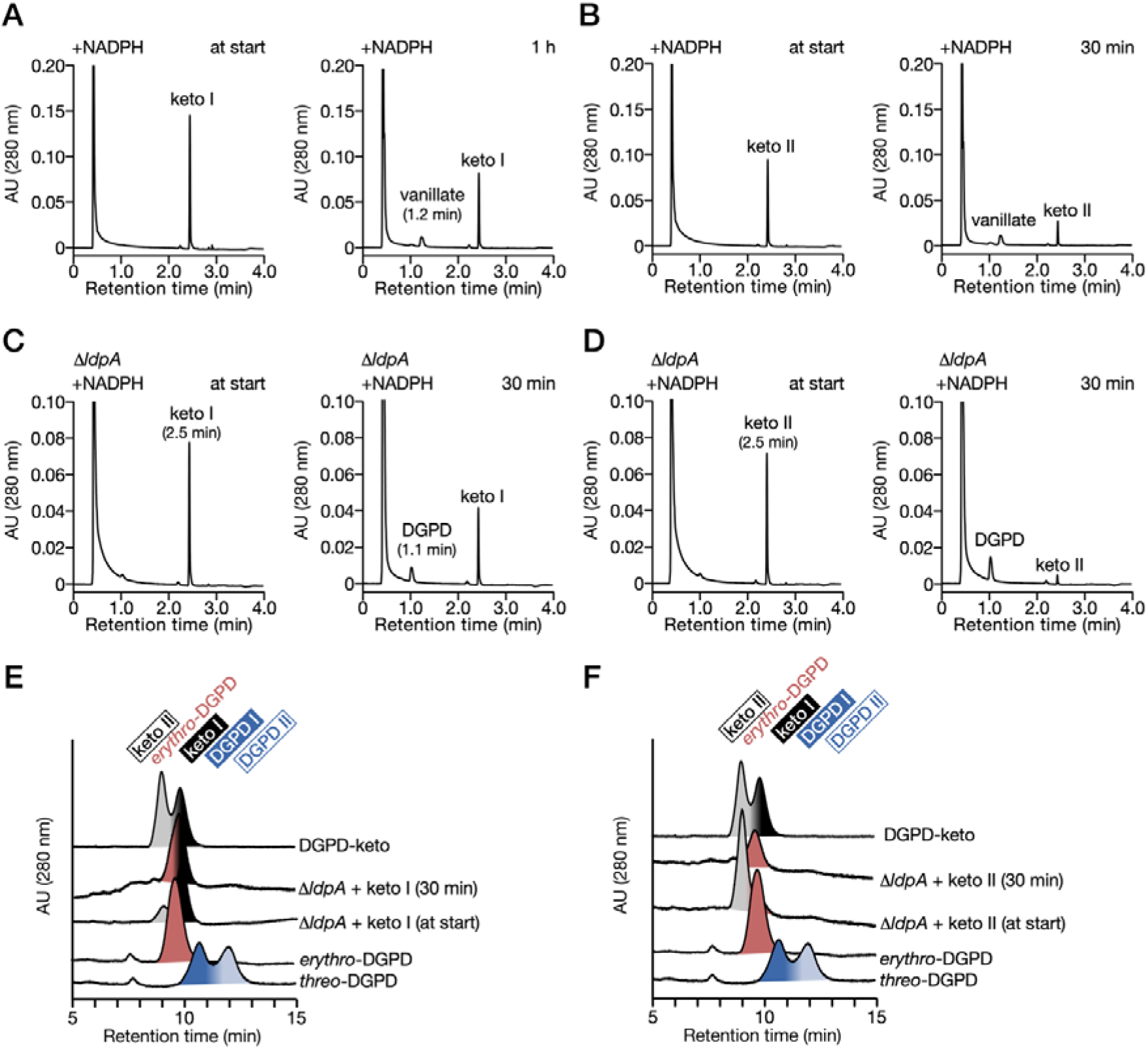
Conversion of DGPD-keto I and II by cell extracts of SYK-6 and *ldpA* mutant. (**A**, **B**) Each of 70 μM DGPD-keto I (**A**) and II (**B**) was reacted with cell extracts of SYK-6 (500 μg/mL) in the presence of 1 mM NADPH for 1 h or 30 min. (**C**, **D**) Each of 70 μM DGPD-keto I (**C**) and II (**D**) were reacted with cell extracts of *ldpA* mutant (Δ*ldpA*, 500 μg/mL) in the presence of 1 mM NADPH for 30 min. (**E**, **F**) Chiral HPLC chromatograms of the products generated in the reactions (30 min) of a Δ*ldpA* cell extract with DGPD-keto I (the same reaction mixture shown in panel **C**) and II (the same reaction mixture shown in panel **D**), respectively.

If ADHs are involved in the formation of *erythro*-DGPD from DGPD-keto using NADPH, the reverse reaction by ADHs may cause the once-generated *erythro*-DGPD to revert to DGPD-keto using NADP^+^ (**Fig. S10A**). Cell extracts of SYK-6 were reacted with *erythro*-DGPD with NAD^+^ or NADP^+^for 2 h, and the reaction products were analyzed by HPLC. As a result, 15% and 30% of *erythro*-DGPD were converted to DGPD-keto in the presence of NAD^+^ and NADP^+^, respectively. In contrast, the remaining 85% and 70% of *erythro*-DGPD were converted to vanillic acid despite the excess NAD^+^/NADP^+^ (1 mM) conditions (**Fig. S10B**). These results suggested that the reaction catalyzed by LdpA will ultimately shift the distribution of the reaction toward further catabolism.

### Identification of the genes responsible for converting DGPD-keto I and II

We further hypothesized that Cα-dehydrogenases are involved in the reduction of DGPD-keto I and II to *erythro*-DGPD and examined their reduction activities toward DGPD-keto I and II. Cell extracts of *E. coli* expressing each of the Cα-dehydrogenase genes except SLG_07240 were reacted with 70 μM DGPD-keto I or II in the presence of NADPH. The activity of the product of SLG_07240, which was not expressed in *E. coli*, was measured similarly using a purified enzyme. However, there were no Cα-dehydrogenases showing DGPD-keto I conversion activity.

In contrast, cell extracts of *E. coli* expressing SLG_12640 converted DGPD-keto II to DGPD (**Fig. 6A**). The specific activity of purified SLG_12640 product toward DGPD-keto II in the presence of NADPH was 62 μmol·min^-1^·mg^-1^, with no activity in the presence of NADH (**Fig. 6B**). No DGPD-keto I conversion activity was observed in the presence of NADH or NADPH. These results indicate that SLG_12640 encodes an NADPH-dependent DGPD-keto II-converting enzyme, and this gene was thus designated *ldpB.* LdpB was also shown to be partially involved in the conversion of DGPD I (**Fig. 4A** and **B**). However, the activity of LdpB toward DGPD I in the presence of NAD^+^ was 0.04% of that toward DGPD-keto II in the presence of NADPH (**Table 1**), and LdpB showed no activity toward *threo*-DGPD in the presence of NADP^+^ (**Fig. S4**). This difference in activity suggests that LdpB essentially acts in the reduction of DGPD-keto II. Also, cell extracts of *E. coli* expressing *ligL* produced a trace amount of DGPD from DGPD-keto II (**Fig. S11**). Since LigL showed weak activity to oxidize *threo*-DGPD to DGPD-keto II in the presence of NADP^+^ (**Fig. S4**), it is likely that DGPD-keto II was converted to DGPD II (*threo*-DGPD) by the reverse reaction.

**FIG. 6.**
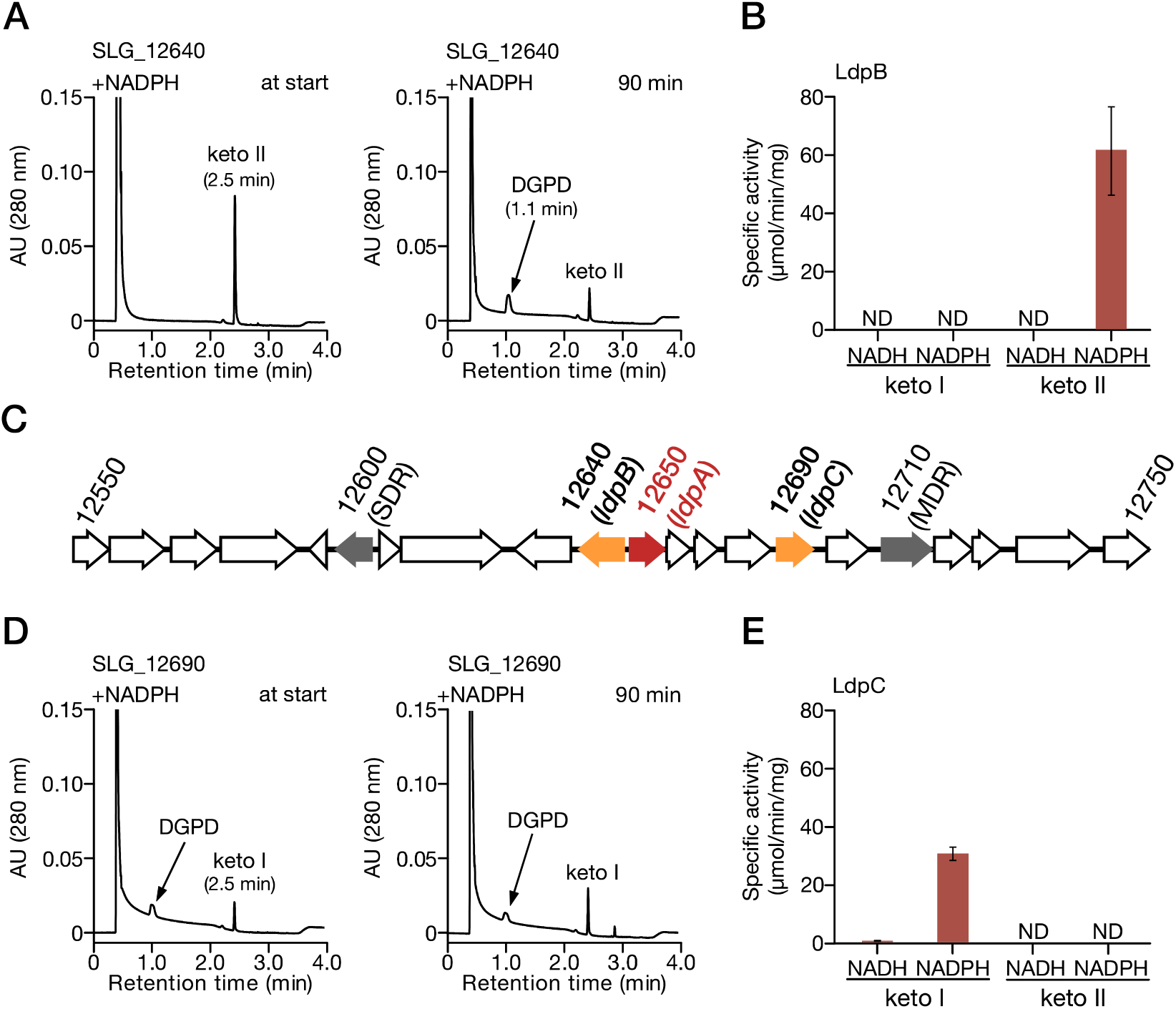
Conversion capabilities of SLG_12640 and SLG_12690 products toward DGPD-keto I and II. (**A**) DGPD-keto II (70 μM) was reacted with a cell extract of *E. coli* BL21(DE3) carrying SLG_12640 (500 μg/mL) in the presence of 1 mM NADPH for 90 min, and then the reaction mixtures were analyzed using HPLC. (**B**) Specific activities of purified SLG_12640 (LdpB) toward DGPD-keto I and II in the presence of 100 μM NADH or NADPH are shown. (**C**) Genetic map around *ldpA*. Two SDR genes, SLG_12600 and SLG_12690, and an MDR gene (SLG_12710) are shown with *ldpA* and *ldpB.* (**D**) DGPD-keto I (70 μM) was reacted with a cell extract of *E. coli* BL21(DE3) carrying SLG_12690 (500 μg/mL) in the presence of 1 mM NADPH for 90 min, and then the reaction mixtures were analyzed using HPLC. (**E**) Specific activities of purified SLG_12690 product (LdpC) toward DGPD-keto I and II in the presence of 100 μM NADH or NADPH are shown. ND, not detected.

We subsequently noted that *ldpB* (SLG_12640) is located directly upstream of *ldpA* (SLG_12650) and that two other SDR genes (SLG_12600 and SLG_12690) and a medium-chain dehydrogenase/reductase (MDR) gene (SLG_12710) are located around *ldpA* (**Fig. 6C**). Among these genes, the two SDR genes were expressed as His-tag fusions in *E. coli*. Cell extracts of *E. coli* expressing each gene (**Fig. S12**) were reacted with DGPD-keto I in the presence of NADPH. HPLC analysis showed that *E. coli* expressing SLG_12690 converted DGPD-keto I to DGPD (**Fig. 6D**), whereas no DGPD-keto I conversion activity was observed in *E. coli* expressing SLG_12600 (**Fig. S13**). The specific activity of the purified SLG_12690 product (**Fig. S12**) toward DGPD-keto I in the presence of NADPH was 31 μmol·min^-1^·mg^-1^, and the specific activity in the presence of NADH was only about 2.4% of that (**Fig. 6E**). No conversion of DGPD-keto II by purified SLG_12690 product was observed in the presence of NADH or NADPH. These results indicate that SLG_12690 encodes an NADPH-dependent DGPD-keto I-converting enzyme, and this gene was designated *ldpC*.

Disruption mutants of *ldpB* and *ldpC* (Δ*ldpB* and Δ*ldpC*) were generated (**Fig. S5**), and their conversion capacities were measured by incubating resting cells of each disruption mutant grown in LB with DGPD-keto I or II. As a result, the amount of DGPD-keto I converted by Δ*ldpC* decreased to approximately 23%of that of the wild type after 4 h of incubation, and Δ*ldpB* almost lost its DGPD-keto II conversion activity (**Fig. 7A** and **B**). Since DGPD-keto I contains approximately 17% DGPD-keto II that could not be removed during preparation, the conversion activity for DGPD-keto I includes the activity toward DGPD-keto II in the sample. Therefore, we performed chiral HPLC analysis on the reaction mixture after incubation of the resting cells of Δ*ldpC* with DGPD-keto I. The DGPD-keto II peak in the sample disappeared, but the DGPD-keto I peak remained almost unchanged (**Fig. 7C**). Thus, the residual activity in Δ*ldpC* was attributed to its activity toward DGPD-keto II. Δ*ldpC* and Δ*ldpB* cells transformed by complementation plasmids regained the ability to convert DGPD-keto I and II, respectively (**Fig. S14**). From these results, it is concluded that LdpC is essentially involved in the conversion of DGPD-keto I and LdpB in the conversion of DGPD-keto II in SYK-6.

**FIG. 7.**
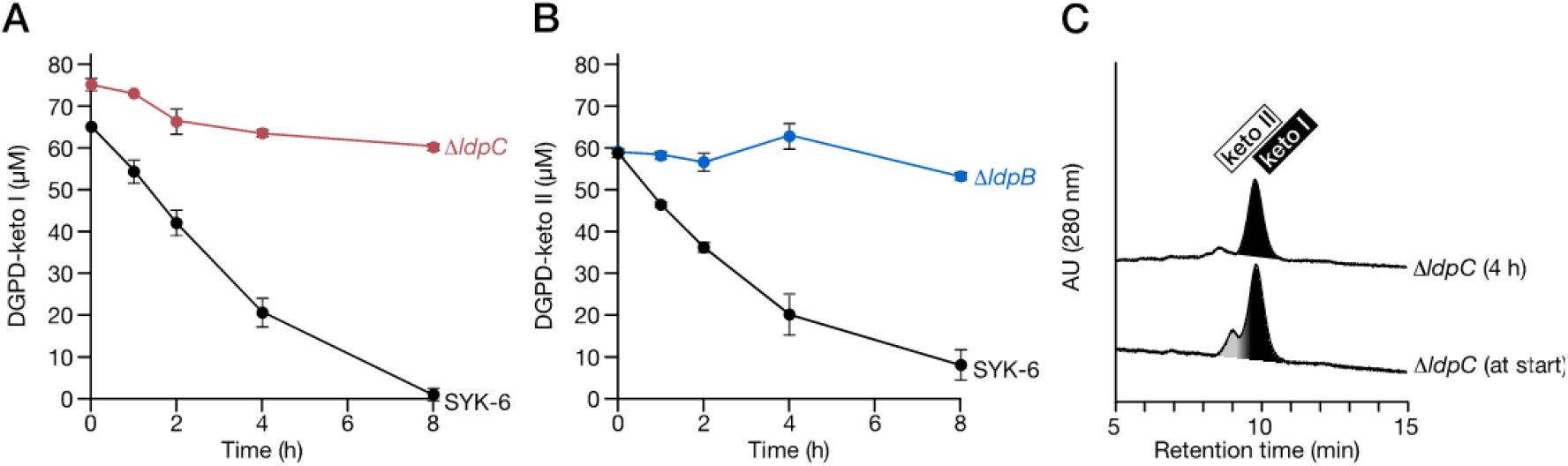
Identification of the enzyme genes involved in the conversion of DGPD-keto I and II in SYK-6. (**A**, **B**) Each of 70 μM DGPD-keto I (**A**) and II (**B**) was incubated with resting cells of Δ*ldpC* and Δ*ldpB* (OD_600_ of 1.0), respectively. Portions of the reaction mixtures were collected over time and analyzed using HPLC. All experiments were performed in triplicate, and each value represents the mean ± standard deviation. (**C**) Chiral HPLC analysis of conversion of DGPD-keto I by resting cells of Δ*ldpC* (OD_600_ of 1.0).

## DISCUSSION

This study revealed that SYK-6 completely catabolizes all stereoisomers of the β-1-type lignin-derived dimer, DGPD, using a stereoinversion mechanism (**Fig. 1**). SYK-6 first stereoselectively oxidizes the Cα alcohol group of *threo*-DGPD to produce DGPD-keto using NAD^+^-dependent LigD and LigL. SYK-6 then stereoselectively reduces the Cα carbonyl group of DGPD-keto using NADPH-dependent LdpB and LdpC to generate *erythro*-DGPD. Finally, the resulting *erythro*-DGPD is converted to DGPD-S by LdpA and undergoes further catabolism. In related work, details of the stereoselective catabolism of GGE, a β-O-4-type lignin-derived dimer, have been elucidated in SYK-6. In GGE catabolism, NAD^+^-dependent LigD and LigL/LigN oxidize the Cα hydroxy group of the four stereoisomers of GGE to α-(2-methoxyphenoxy)-β-hydroxypropiovanillone (MPHPV) enantiomers with a Cα carbonyl group, in a manner analogous to the conversion of *threo*-DGPD to DGPD-keto (30). Then, stereoselective glutathione *S*-transferases (GSTs; LigF and LigE/LigP) cleave the ether bonds of MPHPV by nucleophilic attack of glutathione at the Cβ-position (34, 35). Finally, the resulting glutathione conjugates (α-glutathionyl-β-hydroxypropiovanillone [GS-HPV]) undergo glutathione removal by other GSTs, LigG or LigQ, and are converted to achiral β-hydroxypropiovanillone (36, 37). Thus, the catabolism of β-O-4-type dimer differs from that of β-1-type dimer in that there is a series of enzymes with different stereoselectivity that act on each isomer of substrate and its metabolites (**Fig. 1B; Fig. S2**).

The catabolism of racemic aromatic compounds with stereoinversion is well known; for example, the catabolism of mandelic acid involves a racemase. In *Pseudomonas putida*, *Acinetobacter calcoaceticus*, and *Rhodotorula graminis*, only (*S*)-mandelic acid can be converted to benzoylformic acid by (*S*)-mandelate dehydrogenase to produce benzoylformic acid, which can be further catabolized (38–40). Therefore, (*R*)-mandelic acid is stereoinverted to (*S*)-mandelic acid by mandelate racemase and then further converted. Since mandelate racemase has been shown to be capable of stereoinversion of a wide range of β,γ-unsaturated-α-hydroxy acids, this enzyme has been studied for obtaining their enantiopure compounds (41). Apart from racemase-based deracemization, *in vitro* enzyme systems have been reported that combine two enantiocomplementary ADHs to yield enantiopure compounds of *sec*-alcohols. For example, Hummel and Riebel obtained (*S*)-phenyl ethanol in 100% yield through oxidation of (*R*)-phenyl ethanol and reduction of the resulting keto form by combining an NADPH-dependent (*R*)-phenyl ethanol-specific ADH of *Lactobacillus kefir* and an NADH-dependent (*S*)-phenyl ethanol-specific ADH of *Rhodococcus ruber* DSM 44541 in the presence of both NADP^+^ and NADH (42). It was also shown that the other isomer, (*R*)-phenyl ethanol, could be obtained in 100% yield by switching the enzymes working in oxidation and reduction using NAD^+^ and NADPH as coenzymes. Based on this principle, a set of enzymes has been constructed to stereoselectively obtain *S* or *R* isomers of a wide range of *sec*-alcohols (43).

The DGPD catabolism of SYK-6 appears to have evolved to successfully incorporate the stereoinversion pathway using sequential redox reactions of multiple ADH to convert *threo*-DGPD to *erythro*-DGPD, which undergoes further conversion by LdpA. To our knowledge, such a catabolic strategy has not been reported thus far. There are two main factors that potentially led to the establishment of this reaction system in the same cell. The first point is that the apparent activity of SYK-6 to convert DGPD-keto to *threo*-DGPD (reverse reaction) is weak. DGPD I and II are converted to DGPD-keto I and II in the presence of NAD^+^, mainly by LigD and LigL, respectively (**Fig. 4**). Additionally, the reduction activities of LigD and LigL toward DGPD-keto I and II, respectively, in the presence of NADH, were little or very weak (**Fig. S7**). Moreover, the cell extract of SYK-6 showed no reducing activity to convert DGPD-keto to *threo*-DGPD in the presence of NADH (**Fig. S8**). These findings suggest that DGPD-keto is reduced to *erythro*-DGPD with minimal reverse reaction to *threo*-DGPD, and thus the catabolism can proceed. The second point is that LdpA is active and consumes the product for further conversion in the catabolic pathway. Even under conditions of excess NAD^+^ or NADP^+^, where oxidation reactions are more likely to proceed, cell extracts of SYK-6 produced significantly more vanillic acid than DGPD-keto (**Fig. S10**).

To better understand the ubiquity of the SYK-6-like β-1-type dimer catabolic system, we searched the database using amino acid sequence homology. BLASTP searches were employed using the following parameter, 50% amino acid sequence identity with an e-value of less than 1e-50 to *ldpA* from SYK-6. Homology values are taken directly from BLASTP results. The results showed that *ldpA* orthologs are found in 119 alphaproteobacterial strains (**Table S3**). Of these, 88 strains (approximately 72% of 119 strains carrying *ldpA* orthologs) are in the genera *Novosphingobium* and *Sphingobium.* Next, we searched for strains with *ldpB* and *ldpC* orthologs from strains with *ldpA* orthologs using the same homology criteria. This search revealed that *ldpB* and *ldpC* orthologs are present in 115 and 118 strains, respectively, and 113 strains have both orthologs (approximately 95% of 119 strains carrying *ldpA* orthologs; Group I; **Table S3**). Among the strains belonging to Group I, there are 97 strains that show the same gene arrangement as SYK-6 for *ldpA* and *ldpB*, and 48 strains have *ldpC* in the vicinity of *ldpA* and *ldpB* (within 30 genes). We then searched for strains with *ligD* and *ligL* orthologs in Group I. This search showed that *ligD* and *ligL* orthologs are present in the 44 and 30 strains, respectively. Furthermore, 25 strains are found bearing both orthologs of *ligD* and *ligL* (Group II; **Table S3**). All 25 Group II strains belong to the order Sphingomonad, 24 to the family Sphingomonadaceae, and one to the family Erythrobacteriaceae. Furthermore, 17 strains of Group II are the same genus *Sphingobium* as SYK-6. Compared to strains with orthologs of the catabolism genes of DGPD-keto and *erythro*-DGPD (Group I), fewer strains have orthologs of the catabolism genes of DGPD-keto, *erythro*-DGPD, and *threo*-DGPD (Group II; approx. 22% of Group I). Based on the above, these Sphingomonad strains, including SYK-6, can be regarded as a unique group that can catabolize all β-1-type stereoisomers. Notably, *N. aromaticivorans* DSM 12444, for which the *erythro*-DGPD converting enzyme gene was first identified, was not included in Group II (28). *N. aromaticivorans* DSM 12444 has Saro_1875, which shows 48.7% amino acid sequence identity with SYK-6 *ligL* (e-value, 3e-88), but was excluded from the list due to criteria of >50% identity. SYK-6 cells grown in LB completely degraded 50 μM DGPD I and II in 8 h, whereas *N. aromaticivorans* DSM 12444 converted only DGPD I under the same conditions (**Fig. S15**). Thus, the enzyme gene involved in the conversion of DGPD II in *N. aromaticivorans* DSM 12444 is either nonfunctional under the conditions examined or absent.

Since *ligD* and *ligL* are also involved in GGE catabolism, strains with orthologs of *ligF*, *ligE*, *ligG*, and *ligQ* necessary for GGE catabolism were searched in the Group II strains (30, 34, 37). *ligF* and *ligQ* orthologs were present in all 25 strains (**Table S3**). *ligE* and *ligG* orthologs were present in 24 and 20 strains, respectively. As mentioned earlier, LigG and LigQ are involved in glutathione removal from GS-HPV, with LigG acting only on (β*R*)-GS-HPV and LigQ on both (β*R*)-GS-HPV and (β*S*)-GS-HPV (37). The presence of *ligF*, *ligE*, and *ligQ* orthologs in almost all Group II strains suggests that these strains have both DGPD and GGE catabolic systems. Incidentally, a survey of the database for strains with orthologs of the GGE catabolism genes (*ligD*, *ligL*, *ligF*, *ligE*, and *ligQ*) yielded 24 hits belonging to Group II. These results indicate that both the DGPD and GGE catabolism genes identified in SYK-6 are conserved only in Sphingomonad strains closely related to SYK-6.

Interestingly, Group I also contained *Novosphingobium ovatum* FSY-8, which has three orthologs of *ldpA* (GTZ99_02690, GTZ99_08355, GTZ99_12845). Some of these gene products may possibly convert *threo*-DGPD directly to DGPD-S. Further study will be conducted to this end in the future.

## MATERIALS AND METHODS

### Bacterial strains, plasmids, and culture conditions

The strains and plasmids used in this study are listed in **Table S1**. All strains were cultured in lysogeny broth (LB; 10 g/L of Bacto tryptone, 5 g/L of yeast extract, and 5 g/L of NaCl). SYK-6 and its mutants were cultured in LB containing 12.5 mg/L of nalidixic acid. Complementary strains of SYK-6 and its mutants were cultured in LB containing 50 mg/L of kanamycin (Km) and 1 mM *m*-toluate (an inducer of the expression from P_*m*_ promoter in pJB861) or 12.5 mg/L of tetracycline (Tet) and 100 μM cumate (an inducer of the expression from Q_5_ promoter in pQF). The media for *E. coli* transformants were supplemented with 25 mg/L Km, 12.5 mg/L Tet, 100 mg/L ampicillin (Amp), or 100 mg/L Amp + 25 mg/L chloramphenicol (Cm). SYK-6, its mutants, and complementary strains were cultured at 30°C, and *E. coli* at 37°C, shaking at 160 rpm.

### Preparation of substrates

*threo*-DGPD and *erythro*-DGPD were prepared as described in the supplementary methods or a previous study (44). DGPD-keto was synthesized as shown in the supplementary methods. For the preparation of DGPD I, DGPD II, DGPD-keto I, and DGPD-keto II, racemic *threo*-DGPD (2 mM) was reacted with resting cells (OD_600_ of 10) of *E. coli* BL21(DE3) harboring pET08640 (*ligD*) or *E. coli* harboring pET33660 (*ligL*) in 50 mM Tris-HCl buffer (pH 7.5, buffer A; total volume, 20 mL) at 30°C shaking at 160 rpm for 40 h. Equal volumes of ethyl acetate were added to 20 mL of the reaction solutions, and compounds were extracted three times. The ethyl acetate fractions were concentrated in a centrifugal evaporator until the liquid volumes were reduced to 500 μL. The compounds were separated by thin-layer chromatography (developing solvent, benzene-ethyl acetate-methanol [7:3:2]) and visualized under UV light at 254 nm. An upper spot and a lower spot corresponding to DGPD-keto I and DGPD II from the LigD reaction mixture and an upper spot and a lower spot corresponding to DGPD-keto II and DGPD I from the LigL reaction mixture were cut out, extracted with ethyl acetate three times, finally dissolved in DMSO, and stored at −20°C.

Vanillic acid was purchased from Sigma-Aldrich Co., LLC. NADH, NADPH, NAD^+^, and NADP^+^were purchased from Fujifilm Wako Pure Chemical Corporation.

### Preparation of resting cells and cell extracts

Cells of *E. coli* and SYK-6, its mutants, and complementary strains were grown in LB for 12 h and 24 h, respectively, and were collected by centrifugation (14,000 × *g* for 1 min) and then washed twice with buffer A. The cells were resuspended in the same buffer and used as resting cells. Cells were broken by an ultrasonic disintegrator in a crushed-ice bath at an output ample of 80%, cycle for 10 sec with 1.0-sec intervals every second. This was done three times, and the supernatants of cell lysates were obtained as cell extracts after centrifugation (19,000 × *g* for 15 min). Cell extracts were filtered using an Amicon Ultra centrifugal filter unit (10-kDa cutoff; Merck Millipore) and washed four times with buffer A. The filtrates were used as cell extracts free of molecules below 10 kDa. Protein concentration was determined using the Bradford method with bovine serum albumin as the standard (Bio-Rad Laboratories).

### HPLC and HPLC–MS analysis

HPLC-MS analysis was performed with the ACQUITY UPLC system (Waters) coupled with an ACQUITY TQ detector using a TSKgel ODS-140HTP column (2.1 × 100 mm; Tosoh) as described previously (29, 45) The analytical method for DGPD, DGPD-keto, and vanillic acid was described previously (29). Chiral HPLC analysis was performed with the same HPLC using the CHIRALPAK OZ-H column (4.6 × 250 mm; DAICEL). The mobile phase was acetonitrile containing 0.1%trifluoroacetic acid at a flow rate of 0.5 ml/min. DGPD and DGPD-keto were detected at 280 nm.

### Identification of the metabolites

SYK-6 cell extracts (500 μg or 2 mg protein/mL) were incubated with 100 μM *threo*-DGPD, DGPD-keto I, or DGPD-keto II in the presence or absence of 1 mM NAD^+^, NADP^+^, NADH, or NADPH in buffer A for 1 h or 6 h at 30°C. The reactions were stopped by adding acetonitrile (final concentration, 50%) at the sampling points. Precipitated proteins were removed by centrifugation at 19,000 × *g* for 15 min. The resulting supernatants of the reaction mixtures were analyzed by HPLC, HPLC-MS, and chiral HPLC.

### Expression of SYK-6 SDR genes, including Cα-dehydrogenase genes, and enzyme purification

DNA fragments carrying SLG_07240, SLG_12600, *ldpB* (SLG_12640), *ldpC* (SLG_12690), SLG_28340, and *ligO* (SLG_35880) were amplified through PCR using SYK-6 total DNA and the primer pairs listed in **Table S2**. The amplified fragment of SLG_07240 was cloned into pQF, and other amplified fragments were cloned into pET-16b, respectively, using an NEBuilder HiFi DNA assembly cloning kit. A 1.1 kb NdeI-SalI fragment carrying *ligD* from pETDa, a 1.1 kb NdeI-XhoI fragment carrying *ligL* from pETLa, a 2.8 kb NdeI-EcoRV fragment carrying *ligN* from pETNa were cloned into pET-16b. The nucleotide sequences of the inserts were confirmed by sequencing. pQF07240 was introduced into SYK-6, and other expression plasmids were introduced into *E. coli* BL21(DE3), respectively. The transformed cells were grown in LB. SLG_07240 expression was induced for 24 h at 30°C by adding 100 μM cumate. In the case of *E. coli* transformants, each gene expression was induced for 4 h at 30°C by adding 1 mM isopropyl-β-D-thiogalactopyranoside when the OD_600_ of the cultures reached 0.5. For expression of SLG_28340, *E. coli* harboring pET28340 and pLysS was employed. The cell extracts were prepared as described above. Each protein was purified on a Ni Sepharose High Performance column (GE Healthcare). The purified fractions were desalted and concentrated using an Amicon Ultra centrifugal filter unit (10-kDa cutoff; Merck Millipore), and the enzyme preparations were stored at −80°C. Gene expressions and the purity of the enzymes were examined using SDS-12%polyacrylamide gel electrophoresis. The protein bands in gels were stained with Coomassie Brilliant Blue.

### Stereoselectivity of Cα-dehydrogenases

Purified Cα-dehydrogenases (LigD, LigO, LigN, LdpB, SLG_28340 gene product, and SLG_07240 gene product, 100 μg protein/mL; LigL, 1 μg protein/mL) were reacted with 100 μM *threo*-DGPD in the presence of 1 mM NAD^+^ or NADP^+^ in buffer A for 90 min or 6 h at 30°C. The reactions were stopped by adding acetonitrile (final concentration, 50%) at the sampling time points. The resulting supernatants were analyzed by HPLC and chiral HPLC.

### Activity measurement for reduction of DGPD-keto I and II by purified LigD and LigL

Purified LigD (100 μg of protein/mL) and LigL (1 μg of protein/mL) were reacted with 50 μM DGPD-keto I and II, respectively, in the presence of NADH in buffer A for 90 min or 6 h at 30°C. The reactions were stopped as described above. The resulting supernatants were analyzed by HPLC.

### Activity measurement for reduction of DGPD-keto I and II by SYK-6 SDR family enzymes including Cα-dehydrogenase

Cell extracts of *E. coli* BL21(DE3) harboring pET-16b, pET08640, pET33660, pET35900, pET35880, pET12640, pET12600, pET12690, pET-16b + pLysS, or pET28340 + pLysS (500 μg protein/mL) were reacted with 70 μM DGPD-keto I and II in the presence or absence of 1 mM NADPH for 90 min, respectively. In the case of SLG_07240, a purified enzyme of SLG_07240 gene product (100 μg protein/mL) was used. The reactions were stopped as described above. The resulting supernatants of the reaction mixtures were analyzed by HPLC.

### Activity measurement for reduction of *erythro*-DGPD by SYK-6

Cell extracts of SYK-6 (500 μg of protein/mL) were reacted with 100 μM *erythro*-DGPD in the presence of 1 mM NAD^+^ or NADP^+^ in buffer A for 2 h at 30°C. The reactions were stopped as described above. The resulting supernatants were analyzed by HPLC.

### Activity measurement for oxidation of DGPD I/II and reduction of DGPD-keto I/II by SYK-6 SDR family enzymes, including Cα-dehydrogenases

Purified enzymes (LigD, LigO, and LdpB, 100 μg protein/mL for the conversion of DGPD I; LigL and LigN, 1.0 μg or 100 μg protein/mL for the conversion of DGPD II; LdpC, 1–100 μg protein/mL for the conversion of DGPD-keto I; LdpB, 1-100 μg protein/mL for the conversion of DGPD-keto II) were incubated in 100 μL reaction mixtures containing 50 mM buffer A and 50 μM DGPD I or DGPD II with 500 μM NAD^+^ or NADP^+^ or 50 μM DGPD-keto I or DGPD-keto II with 100 μM NADH or NADPH for 30 sec at 30 °C. Specific activities were calculated by measuring the consumption or production of NADH (ε_340_ = 6400 M^-1^·cm^-1^) and DGPD-keto (ε_340_ = 3200 M^-1^·cm^-1^) or NADPH (ε_340_ = 6400 M^-1^·cm^-1^) and DGPD-keto (ε_340_ = 3200 M^-1^·cm^-1^) using a spectrophotometer (JASCO Corporation). Specific activities of oxidation and reductions were expressed in moles of NADH/NADPH consumed or produced per min per milligram of protein.

### Conversion of DGPD-keto I and II by Δ*ldpA*

Cell extracts of Δ*ldpA* (500 μg protein/mL) were reacted with 70 μM DGPD-keto I and II in the presence of 1 mM NADPH in buffer A for 30 min at 30°C. The reactions were stopped as described above. The resulting supernatants of the reaction mixtures were analyzed by HPLC and chiral HPLC.

### Sequence analysis

Nucleotide sequences were determined by Eurofins Genomics. Sequence analysis and sequence similarity searches were performed using the MacVector program (MacVector, Inc.) and the BLASTP program, respectively (https://blast.ncbi.nlm.nih.gov/Blast.cgi). For phylogenetic analysis, multiple alignments were performed using the MAFFT program (https://www.genome.jp/tools-bin/mafft). Then, a phylogenetic tree was generated using the neighbor-joining algorithm of the MEGA X software, applying 1,000 bootstrap replicates (https://www.megasoftware.net).

### Construction of deletion mutants

To construct the deletion mutants, the upstream and downstream regions of the genes were amplified through PCR from SYK-6 total DNA using the primer pairs listed in **Table S2**. The resulting fragments were cloned into pAK405 or pAK405GFP using an NEBuilder HiFi DNA assembly cloning kit (29, 46). Each resulting plasmid was introduced into SYK-6 cells by triparental mating, and the resulting mutants were selected as described previously (46). Gene deletion was confirmed through colony PCR using the primer pairs listed in **Table S2**.

### Characterization of disruption mutants

Resting cells of SYK-6 and its mutants (for DGPD I conversion, OD_600_ of 10, for conversion of other substrates, OD_600_ of 1.0) were incubated with substrates (DGPD I and II, 50 μM; DGPD-keto I and II, 70 μM) at 30°C with shaking for 8 h or 24 h. For complementation, pQF*ligD*, pQF*ligL*, pQF*ldpB*, and pJB*ldpC* were constructed by cloning the *ligD*, *ligL*, *ldpB*, and *ldpC* fragments amplified by PCR using SYK-6 total DNA and their corresponding primer pairs (**Table S2**) into pQF or pJB861 using an NEBuilder HiFi DNA assembly cloning kit. After confirming the nucleotide sequences of the inserts, each corresponding plasmid was introduced into Δ*ligD*, Δ*ligL*, Δ*ldpB*, or Δ*ldpC* cells through triparental mating. Resting cells of SYK-6 harboring pQF, Δ*ligD* harboring pQF, Δ*ligD* harboring pQF*ligD*, Δ*ligL* harboring pQF, Δ*ligL* harboring pQF*ligL*, Δ*ldpB* harboring pQF, Δ*ldpB* harboring pQF*ldpB*, Δ*ldpC* harboring pJB861, and Δ*ldpC* harboring pJB*ldpC* grown in LB were incubated with substrates as described above. Portions of the cultures were periodically collected, and the amounts of substrates were measured using HPLC.

## ACKNOWLEDGMENTS

This work was partly supported by the Noda Institute for Scientific Research Grant. This work was partially authored by the Alliance for Sustainable Energy, LLC, the manager and operator of the National Renewable Energy Laboratory for the U.S. Department of Energy (DOE), under Contract No. DE-AC36-08GO28308. EK and GTB were funded by The Center for Bioenergy Innovation, a U.S. DOE Bioenergy Research Center supported by the Office of Biological and Environmental Research in the DOE Office of Science. RK and GTB thank the U.S. Department of Energy Office of Energy Efficiency and Renewable Energy Bioenergy Technologies Office for funding. The views expressed herein do not necessarily represent the views of the DOE or the U.S. Government. The U.S. Government retains and the publisher, by accepting the article for publication, acknowledges that the U.S. Government retains a non-exclusive, paid-up, irrevocable, worldwide license to publish or reproduce the published form of this work, or allow others to do so, for U.S. Government purposes.

Conceptualization, GTB, NK, and EM; Validation, RKato; Investigation, RKato, KM, SK, SH, RKatahira, MN, YH, and EK; Writing – original draft, RKato, SH, NK, and EM; Writing – reviewing and editing, RKato SH, RKatahira, YH, EK, GTB, NK, and EM; Supervision, GTB, NK, and EM; Project Administration, GTB and EM; Funding acquisition, GTB and EM.

